# Leveraging Targeted Gene Sets and Neural Networks for Zebrafish Transcriptome Extrapolation in High-Throughput Toxicogenomics

**DOI:** 10.64898/2026.08.07.743325

**Authors:** BE Howard, D Mav, M Balik-Meisner, D Phadke, EH Scholl, AJ Green, L Truong, RL Tanguay, RR Shah

## Abstract

**Background:** Zebrafish (Danio rerio) are a powerful vertebrate model for developmental toxicology and chemical safety assessment, yet large-scale transcriptomics in zebrafish remains limited by cost and data heterogeneity. Targeted transcriptomics offers a cost-effective alternative, but gene extrapolation methods tailored to zebrafish have not been systematically developed or evaluated.

**Objectives:** While the S1500+ platform is widely used for toxicogenomics research with rat, mouse, and human cell lines as model systems, its use in zebrafish has been limited due to data scarcity and lack of suitable bioinformatics approaches for analysis of such data. To that end, we sought to (i) curate a large zebrafish transcriptomic training data resource, and (ii) evaluate multiple machine learning strategies for reconstructing unmeasured transcriptome-wide expression profiles for data originating from the zebrafish-specific reduced representation gene set (“Zf S1500+”).

**Methods:** We assembled 14,924 zebrafish RNA-Seq samples covering 21,930 genes across 1,246 studies. Using the Zf S1500+ gene subset (3,062 genes), we trained and tested three extrapolation approaches: principal components regression (PCR), a locally weighted extension of PCR (PCR+), and a neural network mixture-of-experts model (NN-MoE). Model performance was assessed using mean absolute error (MAE), mean squared regression error (MSRE), and weighted variants of these metrics.

**Results:** Extrapolation performance using the baseline approach was strongly influenced by tissue and developmental context, with within-tissue models outperforming cross-tissue models. Errors were lowest when training and testing were conducted within the same tissue or between developmentally related tissues. Both PCR+ and NN-MoE improved upon the baseline PCR approach, with NN-MoE reducing average MAE by ∼20% and MSRE by ∼17%. Importantly, extrapolation remained reliable for the majority of genes, even when limiting output to high-confidence predictions using an empirical MAE threshold.

**Conclusions:** We demonstrate that targeted transcriptomics can be effectively extended to zebrafish, enabling robust transcriptome-wide extrapolation at reduced cost. The NN-MoE method provided the most substantial gains, highlighting the value of non-linear and ensemble modeling in heterogeneous datasets. These results establish a scalable framework for zebrafish toxicogenomics and suggest that accuracy will continue to improve with larger, better-annotated datasets, paving the way for broader application in chemical safety assessments.

## Introduction

The zebrafish (*Danio rerio*) has become an invaluable model in biomedical and toxicological research due to its well-annotated genome, genetic similarity to humans, rapid development, optical transparency during early life stages, and amenability to large-scale *in vivo* screening [1], [2], [3]. These attributes make zebrafish particularly attractive for developmental toxicology studies, where high-throughput systems are increasingly needed to evaluate the vast number of environmental chemicals with potential human health relevance.

High-throughput transcriptomics (HTTr) has emerged as a transformative approach for characterizing chemical effects on biological systems [4]. By capturing broad patterns of gene expression, HTTr provides mechanistic insights into chemical activity and enables scalable hazard screening. However, the cost of whole-transcriptome measurement remains a limiting factor when applied at the scale required to assess thousands of chemicals.

Targeted platforms such as S1500+ and L1000 have been developed to address this challenge [5], [6], [7], [8]. These assays measure a carefully selected subset of genes at substantially reduced cost. Because gene expression patterns are highly correlated, this reduced representation can be leveraged to infer the expression of unmeasured genes. Building on this principle, Sciome, in collaboration with NIH and EPA, previously developed a principal components regression (PCR) method that uses the S1500+ gene set measured on the TempO-Seq platform to estimate the unobserved transcriptome [8]. This approach is now routinely employed by NIH, EPA, and other groups for high-throughput chemical assessment [9], [10], [11], [12].

Compared to other organisms such as mice and humans, the total number of samples currently available for training a gene extrapolation method for zebrafish is reduced by more than an order of magnitude. In addition, there exists a high degree of heterogeneity in zebrafish samples, which can encompass multiple strains, as well as diverse tissue types, developmental stages, and environmental exposures. These features motivated us to further investigate alternative extrapolation approaches alongside our baseline method to ensure that we make the most efficient use of the available training data while maintaining optimal performance across a diverse range of populations and phenotypes.

In the present work, we extend our targeted extrapolation strategy to zebrafish within the context of developmental toxicology screening. Specifically, we:

1. Curate a comprehensive zebrafish dataset comprising 14,924 transcriptome samples, each with 21,930 measured genes.
2. Evaluate and validate three approaches for transcriptome extrapolation—standard PCR, an enhanced PCR+ variant, and a neural network “mixture of experts” (NN-MOE) method—highlighting conditions under which each performs optimally.
3. Demonstrate that extrapolated expression measurements can increase the coverage of key biological pathways while enhancing the signal-to-noise ratio in the observed differential transcriptomic changes.

Together, these contributions establish a robust framework for cost-efficient transcriptome extrapolation in zebrafish, thereby facilitating scalable application of developmental toxicogenomics in chemical safety assessment.

## Methods

### Zf S1500+ Gene Set

The zebrafish S1500+ gene set was developed using a combination of zebrafish orthologs for human S1500+ genes and gene nominations from developmental neurotoxicity (DNT) and zebrafish field experts [7]. The most recent version of the Zf S1500+ gene set contains 3,062 genes. In the original publication, we examined the feasibility of extrapolating the full zebrafish transcriptome from this subset of genes to make pathway-level inferences. While this approach led to biologically meaningful results, the authors suggested that extrapolation accuracy could improve as the quantity of publicly available transcriptomic zebrafish data increases.

### Data Acquisition, Preparation, and Processing

We obtained a large collection of publicly available Zebrafish RNA-Seq data from the NCBI Sequence Read Archive (SRA). By querying the SRA metadata for *Danio rerio* RNASeq samples from a transcriptomic library source sequenced on a high-throughput Illumina platform, we were able to identify an initial set of 36,069 SRA runs from 28,985 BioSamples.

Fastq files for the individual runs were downloaded and processed by a pipeline that used FastQC [13] to collect sequence and quality metrics. Count matrices at the transcript level were created via pseudo alignment using Kallisto v0.51.1 [14] and an index created from the CDS file for NCBI’s zebrafish genome assembly (GRCz11). These counts were then aggregated into gene-level counts by totaling the transcripts associated with each gene. For BioSamples containing more than one run, we totaled read counts across the individual runs.

We required a minimum of 5 million pseudo-aligned reads for each BioSample, with an alignment rate of at least 25%. Furthermore, at least 35% of all genes and 45% of the Zf S1500+ genes were required to be expressed with at least one assigned read. To ensure that a small number of genes were not capturing the majority of the reads in a BioSample, we required that 90% of the total counts were captured by no less than 8% of the genes.

To eliminate single-cell RNA sequencing samples, we downloaded the GEO Metadata for those samples that have GEO IDs. Because NCBI does not have a specific field to indicate single-cell samples, we heuristically examined the metadata for indications of scRNA sequence, including phrases like “InDrop” or “SMARTSeq” and filtered matching samples from the final dataset.

After applying the above filters, we were left with 14,924 high-quality full-transcriptome samples for model development, a nearly 10-fold increase compared to our previous publication. TPKM normalization was performed separately for genes that occur in the Zf S1500+ subset and for genes that do not occur in the Zf S1500+ subset. TPKM normalized counts where subjected to log2 transformation (post addition by 1 to avoid negatives).

### Three Algorithms for Extrapolation

#### Principal Components Regression (PCR)

The principal components regression (PCR) approach for transcriptome extrapolation was previously described in detail in our 2018 publication [8]. Briefly, the principal components are first computed from expression values of the Zf S1500+ genes in a training data set (denoted *X*). Using these components, a least squares regression is performed to model the expression of unobserved genes (*Y*) based on the observed Zf S1500+ values. For new datasets, where Zf S1500+ measurements (*X*) are available but *Y* values are unmeasured, the observed data are projected onto the principal components derived from the training set. Predicted expression values (*Ŷ*) for the unmeasured genes are then obtained by applying the regression weights estimated from the model.

#### Locally Weighted Principal Components Regression (PCR+)

In order to address the heterogeneity of the training data, we propose an extension to our baseline PCR approach. The modified approach (PCR+) builds on the PCR method, except instead of simple least squares linear regression, we construct a new regression model “on-the-fly” for each test set, weighting training samples by their distance from the observed test set *X* values. In this way, samples that are similar to the dataset being extrapolated are given more weight when estimating regression weights to predict the unobserved genes.

For the distance score, we used Euclidean distance in principal component space. Measured genes were projected into a common coordinate system as defined by the principal components of the full training set. Gene expression values were averaged over all samples in the test set. The distance for each training sample was computed relative to the test sample coordinates.

#### Neural Network Mixture of Experts (NN-MoE)

Motivated by the desire to better model data heterogeneity and also to accommodate non-linear regression relationships, we evaluated a novel neural network approach to gene extrapolation. The architecture we employ (**Figure 1**) is a variant of the “Mixture of Experts” ensemble approach which has recently gained popularity in Large Language Models including Mixtral and DeepSeek R1 [15], [16]. Our model consists of a set of *N* fully connected feedforward networks, each of which takes as input the measured values for the Zf S1500+ genes. Each of these “regression heads” is designed to independently predict the expression values for the unmeasured genes. In parallel, the observed Zf S1500+ genes are also passed into a separate classification network which is trained to predict which of the regression heads is best suited for estimating the unobserved genes in a given sample. In each training step, the classification error is computed as the cross-entropy loss between the raw (not soft-maxed) classification predictions and a one-hot representation of the performance of the *N* regression heads, with the preferred choice being the regression head that minimizes the MAE on the training sample. The output prediction for the model is a weighted average of the individual regressor head predictions, with weights taken from the soft-maxed output of the classification model. The total loss at each round of training is an average of the regression mean squared error and the classification cross-entropy loss. For the zebrafish data set we used *N* = 10 regression heads. The model was trained for 500 epochs using the AdamW optimizer with a learning rate of .00045 and the pytorch cosine learning rate scheduler with warmup. We were able to train the model in less than 4 hours using an RTX 4090 GPU.

**Figure 1:**
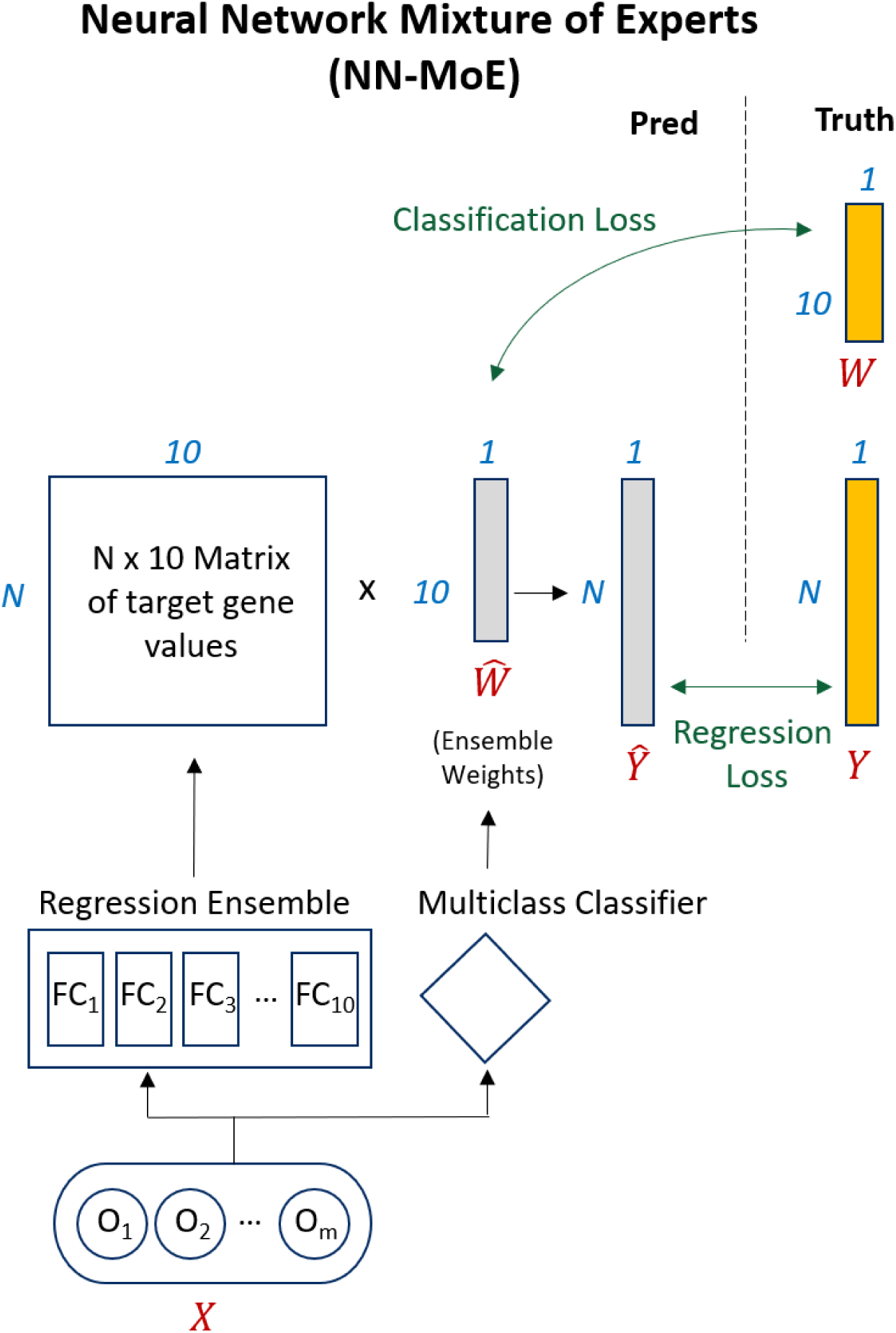
Schematic representation of NN-MoE network. The method utilizes *N* independent neural regression heads with a multiclass classifier to assign weights to training samples.

#### Evaluation Criteria and Experimental Approach

To train and evaluate extrapolation models, a subset of the available data was randomly withheld as a test set, while the remaining data were used for model training. The test set included randomly selected studies as well as 148 RNA-seq samples from a recent developmental neurotoxicity (DNT) transcriptomics experiment [17]. This study (hereafter referred to as the “DNT Test Study”) was designed to characterize transcriptional changes in whole 48 hpf zebrafish following exposure to a panel of 27 DNT chemicals at concentrations associated with morphological phenotypes observed in 120 hpf larvae.

The complete test set comprised approximately 891 out of the 14,924 available samples. Because samples within a given GEO series typically include replicates and are generally more similar to each other than to samples from unrelated series, care was taken to ensure that all samples from the same GEO series were assigned to the same subset (training or test). When training the neural network, a small validation subset was also set aside from the training data to periodically evaluate the performance of the model.

After training models using each of the three extrapolation methods and the training data subset, accuracy of the extrapolated values was evaluated on the test set using the following metrics: MAE, MSRE, weighted-MAE and weighted-MSRE. The MAE (median absolute error) is simply an average of the absolute difference between the estimate and true value for each gene expression level, averaged over all predicted genes. The MSRE (mean squared regression error) is the mean of the squared differences, again averaged over all predicted genes. The MSRE tends to penalize genes with large errors more harshly, while the MAE can be more robust to outliers. The weighted versions of each of these metrics gives increased emphasis to genes that are more highly expressed, weighting the contribution from each gene by its overall expression across all samples. Since each of the four metrics is a measure of error, lower values indicate improved performance.

To better understand the impact of gene extrapolation on downstream biological interpretation, we performed a detailed analysis of the results from the DNT Test Study. Specifically, we performed a comprehensive differential gene expression analysis and pathway enrichment analysis using (1) the genes from the measured Zf S1500+ genes only, (2) the measured Zf S1500+ genes plus the extrapolated non-S1500+ values from each of the extrapolation methods, and (3) the complete RNA-seq dataset which includes measured expression values for both Zf S1500+ and non-S1500+ genes. We utilized Student’s t-test statistics to measure gene-level differential activity and identify differentially expressed genes (DEGs). The significance p-value for differential expression was computed using a permutation approach utilizing 10,000 random permutations of treatment labels. For a gene to be considered significantly differentially expressed, it was required to have an absolute foldchange ≥ 1.5 and permutation test p-value ≤ 0.005. DEG accuracy was evaluated using the following metrics: F1 score, sensitivity, and precision.

Subsequently, significant DEGs were used as input for pathway analysis. Fisher’s exact test was used to obtain differentially enriched pathways (DEPs) that were significantly enriched with DEGs. Pathway analysis was performed on all Canonical Pathways (C2-CP), KEGG, REACTOME, WIki, BIOCARTA, and Hallmark pathways from the Molecular Signature Database (MSigDB version 7.5.1) having five or more human genes orthologous to zebrafish genes present the GRCz11 transcriptome definition. For a pathway to be considered as a significantly differentially enriched pathway, it was required to have a Benjamini & Hochberg adjusted p-value ≤ 0.05.

## Results

### RNA-Seq Dataset Characteristics

The full RNA-Seq dataset contains 14,924 samples from 1,246 distinct studies measuring the expression of 21,930 genes. Expression values were captured in a 14,924 x 21,930 “signal matrix” which contains normalized read counts (log_2_( TPKM + 1)) for each gene measured in those samples. To get a sense of the average, range and variability of the expression values, the mean and variance were computed for each gene across samples. The 50th percentile (median) for mean gene expression occurred at 3.1 and the median gene variance across samples was 1.6.

All genes were expressed in at least one sample. The median percentage of genes expressed in the studies was 91.83% and the smallest percentage of expressed genes in any study was 38.11%. The mean correlation of gene expression across samples was 0.08 with a standard deviation of 0.23. The median gene to gene correlation (excluding the diagonal of the correlation matrix) was 0.06. The minimum correlation was −0.85 and the max was 1.0. On average, each gene has maximum correlation of 0.75 with some other gene.

The mean number of samples per study is 12. Most studies (826/1246 or 66.3%) have 10 or fewer samples, but 6 studies (0.5%) contain more than 100 samples. Half of the samples come from just 183 studies (14.7%).

Using the available meta data extracted from the GEO database, we identified 281 distinct tissue label annotations, 27 distinct strain labels, 6 distinct dev stage labels. The top 25 tissue labels represent 73.9% of the samples.

### Zebrafish Extrapolation Methods Comparison

**Table 1** shows values of performance metrics averaged across all of the extrapolated genes for each of the three methods. Both PCR+ and NN-MoE exhibit improved performance compared to the baseline (PCR). For example, compared to the baseline, NN-MoE reduced the mean MAE by 18.3% and the mean MSRE by 14.1%, on average, while the PCR+ method reduced the mean MAE by 9.7% and the mean MSRE by 7.3%, on average.

**Table 1:** Extrapolation performance of three algorithms.

| Method | All Genes (18,868) |  |  |  | High Confidence Estimates<br>(approx. 90% of all genes) |  |
| --- | --- | --- | --- | --- | --- | --- |
|  | MAE | Weighted<br>MAE | MSRE | Weighted<br>MSRE | MAE | MSRE |
| **PCR**<br>(baseline) | 0.2139 | 0.2246 | 0.2201 | 0.2254 | 0.1974 | 0.1897 |
| **PCR+** | 0.1931 | 0.2014 | 0.2041 | 0.2076 | 0.1772 | 0.1744 |
| **NN-MoE** | 0.1748 | 0.1890 | 0.1890 | 0.1860 | 0.1583 | 0.1584 |

Regardless of the algorithm, there are a small number of genes with high extrapolation error rates. The genes with the highest NN-MoE error rates (MAE) are shown in **Table 2**, and a cumulative plot of the NN-MoE MAE and MSRE is shown in **Figure 2**.

**Figure 2.**
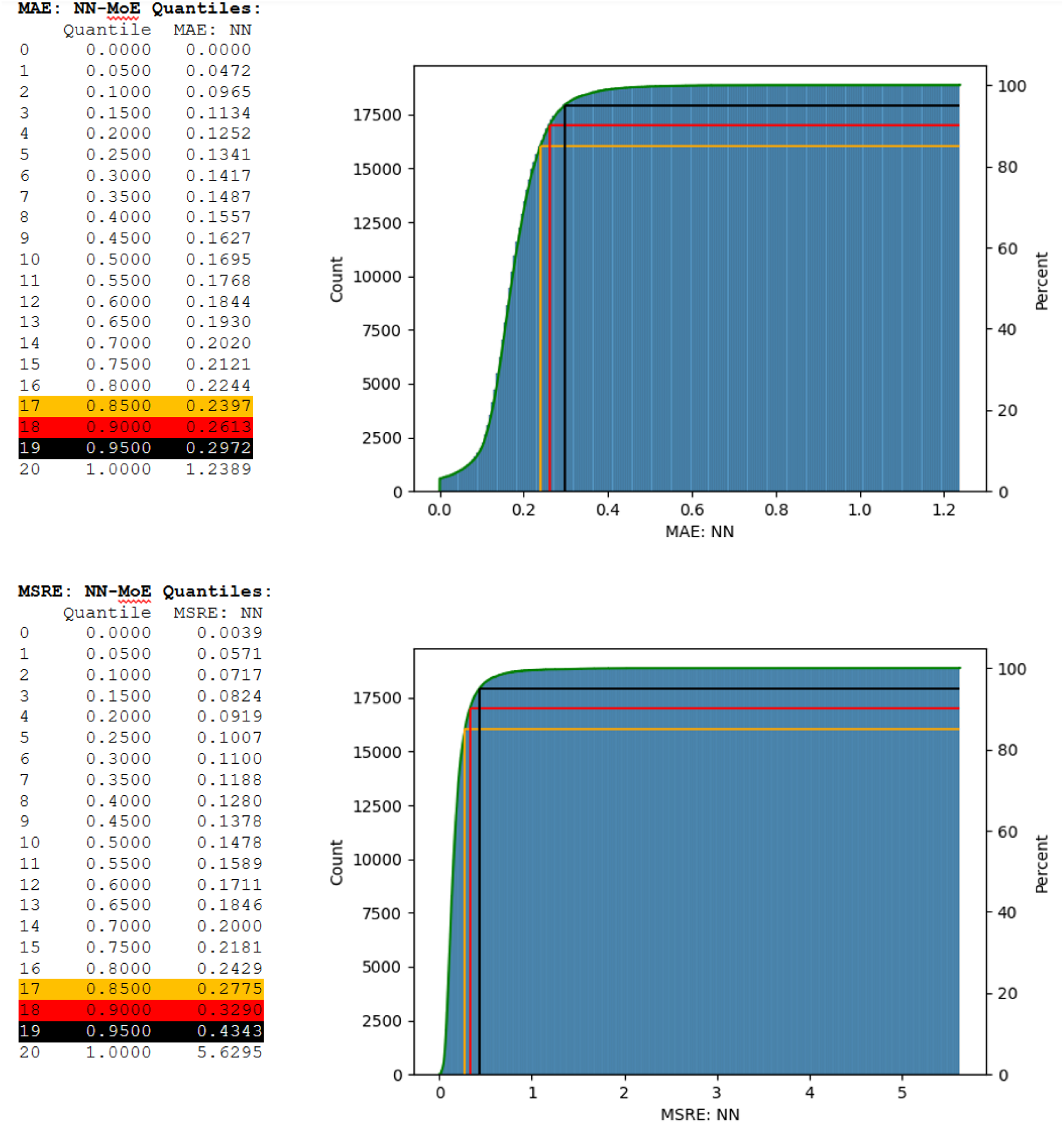
Distribution of MAE and MSRE scores for NN-MoE network.

**Table 2:** 10 genes with highest extrapolation error using NN-MoE.

| GeneID | Symbol | Description | MAE: PCR | MAE: PCR+ | MAE: NN |
| --- | --- | --- | --- | --- | --- |
| 337770 | actc1c | actin, alpha,<br>cardiac muscle 1c | 1.4165 | 1.4243 | 1.2389 |
| 100149863 | her4.1 | hairy-related 4,<br>tandem duplicate 1 | 1.2225 | 0.9929 | 0.9420 |
| 564515 | si:ch211-121a2.2 | si:ch211-121a2.2 | 0.8053 | 0.7492 | 0.8109 |
| 494086 | pih1d2 | PIH1 domain<br>containing 2 | 0.8045 | 0.8524 | 0.7749 |
| 100535646 | si:ch211-213a13.1 | si:ch211-213a13.1 | 0.7814 | 0.8142 | 0.7625 |
| 100536270 | lsm10 | LSM10, U7 small<br>nuclear RNA<br>associated | 0.5954 | 0.6870 | 0.7516 |
| 100034445 | znf1179 | zinc finger protein<br>1179 | 0.9165 | 0.7992 | 0.7515 |
| 791449 | zgc:113363 | zgc:113363 | 0.8187 | 0.7282 | 0.7373 |
| 797669 | ccl39.6 | chemokine (C-C motif) ligand 39, duplicate 6 | 0.7545 | 0.8510 | 0.6999 |
| 100148329 | her4.2 | hairy-related 4, tandem duplicate 2 | 0.632785275 | 0.653738145 | 0.6990 |

Note that by setting a threshold of, for example, 0.27 expected MAE, we can still make predictions for the majority of the unmeasured genes, in this case 90% of the transcriptome. After applying this threshold, the relative overall performance improvements observed with the two alternative extrapolation algorithms is similar to that observed when attempting to predict the entire transcriptome. For example, compared to the baseline, NN-MoE reduced the mean MAE by 19.78% and the mean MSRE by 16.5%, on average, while the PCR+ method reduced the mean MAE by 10.21% and the mean MSRE by 8.1%, on average.

### Impact of Data Heterogeneity on Extrapolation Performance

To examine the impact of training data heterogeneity on extrapolation performance, particularly in the context of small datasets, we annotated samples in our dataset according to tissue type. Tissue terms were derived from the GEO [18] *CHARACTERISTICS_CH1* sample metadata field. However, this metadata is often inconsistent: it lacks a controlled vocabulary and is frequently incomplete or inaccurate. To address these limitations, we applied a heuristic mapping strategy to align the raw GEO tissue labels with controlled vocabularies from the Zebrafish Anatomy and Development Ontology [19] and the BRENDA Tissue and Enzyme Source Ontology [20]. Each GEO-derived term was matched to ontology terms using a composite scoring system that combined lexical similarity (Levenshtein distance) with semantic similarity derived from Specter language model embeddings [21].

Next, we restricted our analysis to tissues represented by at least 150 samples in the dataset. Samples lacking tissue annotations, as well as those labeled with ambiguous terms (e.g., developmental stages such as “embryo” or “larva” or generic anatomical regions such as “head” or “whole organism”), were excluded. This filtering step yielded a set of eight tissues for analysis. For these tissues, we systematically evaluated all 8 × 8 possible training–testing combinations. Specifically, for each pairing of Tissue A (training tissue) and Tissue B (test tissue), we identified a study containing at least 30 samples for Tissue B (hereafter “Test Study B”). We then assessed whether the remaining studies provided at least 100 independent samples for Tissue A (hereafter “Tissue A Samples”). If these conditions were met, we constructed a training set (“Training Dataset A”) by randomly selecting 100 samples from Tissue A Samples, and a test set (“Test Dataset B”) by randomly selecting 30 samples from Test Study B. Importantly, the training and test sets were always drawn from distinct studies, even when the training and testing tissues were the same.

For each train/test pair, we fit a baseline PCR extrapolation model to Training Dataset A and evaluated its performance on Test Dataset B. Based on results from individual tissue comparisons (e.g., brain vs. liver), we hypothesized that models trained and tested on the same tissue would exhibit lower error, as measured by mean absolute error (MAE) and mean squared relative error (MSRE), than models trained and tested on different tissues.

**Figure 3** summarizes the results in a matrix where each cell represents the weighted MAE score for a given train/test tissue pair. Rows indicate the training tissue, and columns indicate the testing tissue. To enable comparison across tissues, scores in each row were normalized by the corresponding diagonal value (i.e., the unnormalized MAE from training and testing on the same tissue). Thus, diagonal values are fixed at 1. Scores greater than 1, shown in red, indicate that extrapolation performance was worse than the within-tissue baseline for that test tissue. White colored cells (including diagonals) represent values near 1, indicating performance comparable to the baseline. Green cells highlight cases where models trained on one tissue generalized better to a different tissue than to the same tissue, yielding normalized scores below 1.

**Figure 3:**
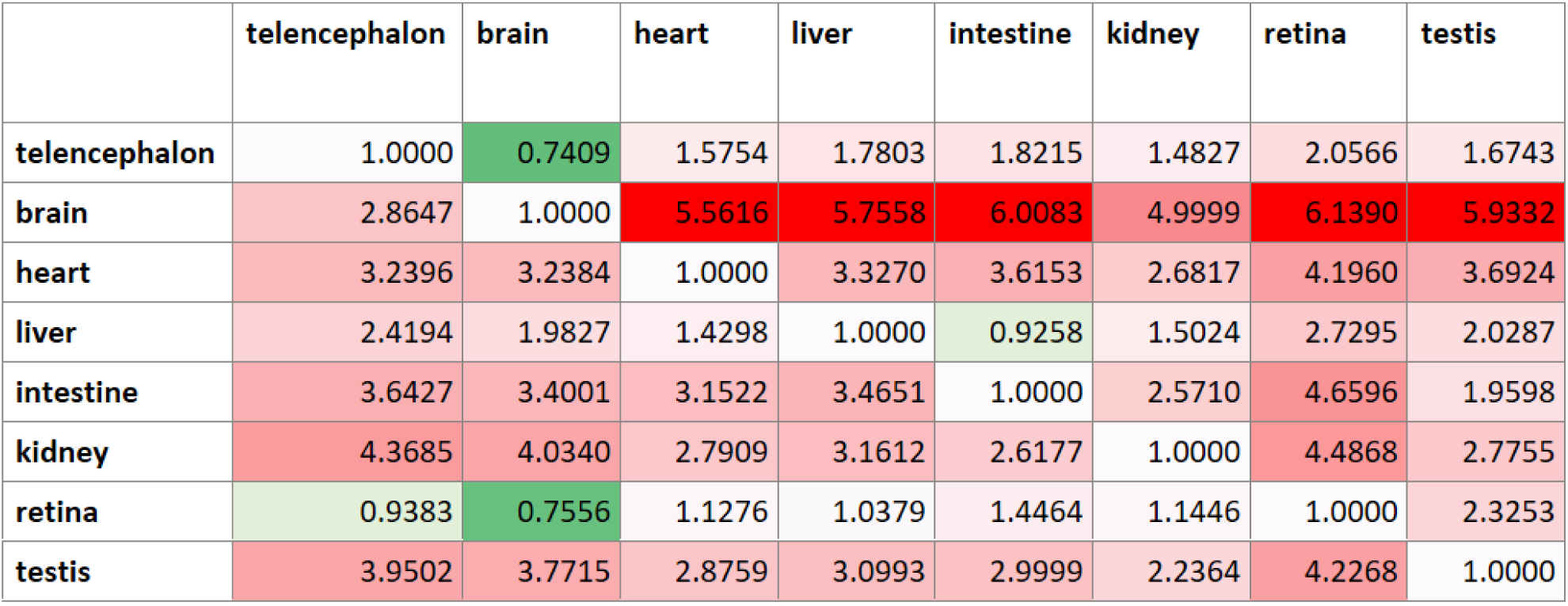
Impact of tissue type mismatch on extrapolation results.

The matrix is dominated by red cells, indicating frequent decreases in extrapolation performance when training and testing are conducted across different tissues. In contrast, the rare off-diagonal white and green cells, which represent performance comparable to or better than the within-tissue baseline, tend to occur between tissues that are anatomically related (e.g., brain and telencephalon) or that share common developmental origins. Specifically, tissues derived from the same embryonic germ layer often show greater cross-tissue generalizability (Ectoderm/neuroectoderm: telencephalon, brain, retina; Mesoderm: heart, kidney, testis; Endoderm: liver, intestine).

### NN-MoE Ensemble Weights

**Figure 4** shows the learned NN-MoE ensemble weights for several studies in the data set after training. The weights for the 10 regression heads are shown in the numbered columns. For these studies, the regression heads number 1 and number 9 appear to be differentially activated according to “tissue type” and other latent meta data associated with individual studies. The remaining regression heads are also weighted differently in various studies, though the change is not as dramatic. This suggests that the NN-Moe approach has learned to implicitly model data heterogeneity using an ensemble of specialized expert predictors.

**Figure 4.**
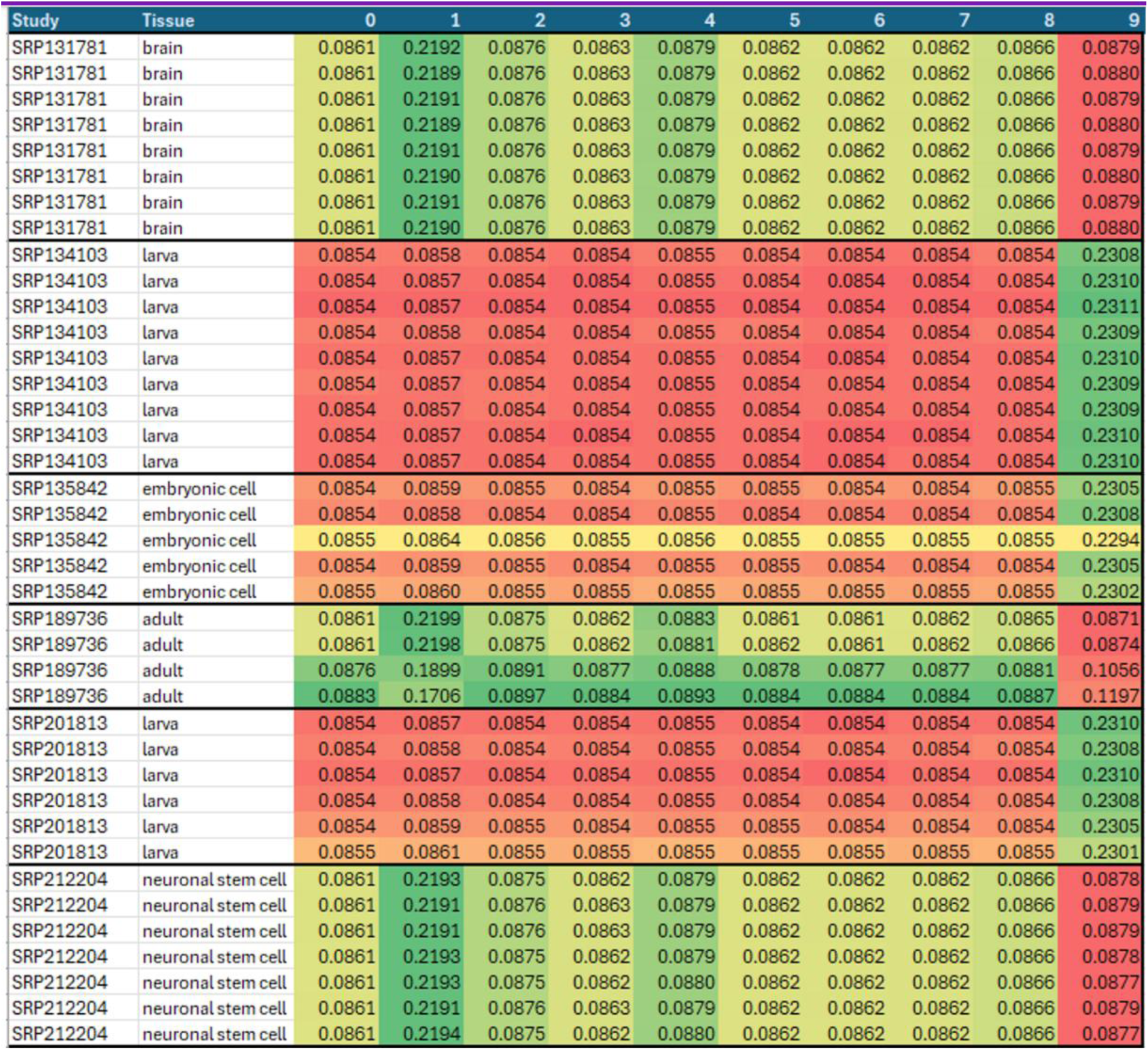
Examples of NN-MoE regression head weights for heads 0 – 9. Note that regression head 9 seems to upweighted for larval and embryonic samples, while regression head 1 is upweighted for neural and brain samples.

### Zebrafish Pathway Coverage

There are 46 zebrafish WikiPathways [22], of which 45 contain at least 5 genes. **Table 3** illustrates the potential improvement in pathway coverage when performing genome extrapolation on the genes measured by the Zf S1500+ platform. When limiting the analysis to only the measured genes, 5 of the 46 pathways were missed completely. Overall mean pathway coverage was 40.7% of the pathway genes. By including the full extrapolated transcriptome (21,930 genes), pathway representation increased to 45 out of 46 pathways, with a mean coverage of 99.5% of the genes in each pathway. Furthermore, even when stringent criteria are used to exclude the small subset of genes having high prediction error on the training set, pathway coverage remains greatly enhanced compared to using only the measured genes.

**Table 3:** Zebrafish WikiPathway coverage for measured and extrapolated genes.

| Platform | # WikiPathways with $\geq 5$ genes | Mean Pathway Coverage |
| --- | --- | --- |
| S1500+ | 41 | 40.7% |
| Whole Extrapolated Transcriptome (21,930 genes) | 45 | 99.5% |
| Whole Extrapolated Transcriptome excluding genes with MAE>0.30 for both methods (21,145 genes) | 45 | 96.7% |
| Whole Extrapolated Transcriptome excluding genes with MAE>0.25 for both methods (19,516 genes) | 44 | 91.0% |
| Whole Extrapolated Transcriptome excluding genes with MAE>0.20 for both methods (14,486 genes) | 43 | 77.8% |

### Potential For Further Improvement with Additional Training Data

**Figure 5** shows the relative performance of the three extrapolation algorithms for zebrafish alongside 3 additional organisms. While not part of the current study, Sciome has previously curated these datasets for use with our baseline PCR extrapolation model and they are useful here for the purposes of comparison. The results are arranged from top to bottom in order of increasing training data availability. For zebrafish, our latest dataset contains a total of 14,924 samples while our previously curated datasets for rat, mouse, and human range in size from 26,716 to 463,055 samples. Viewing the graphs from top to bottom, there is a visible rightward shift indicating that the performance of the PCR+ and NN-MoE models continue to improve relative to the baseline (PCR) with increased training data. In addition, the median MAE and MSRE for NN-MoE extrapolated genes tends to continue to decrease with increased training data availability (**Table 4**).

**Figure 5:**
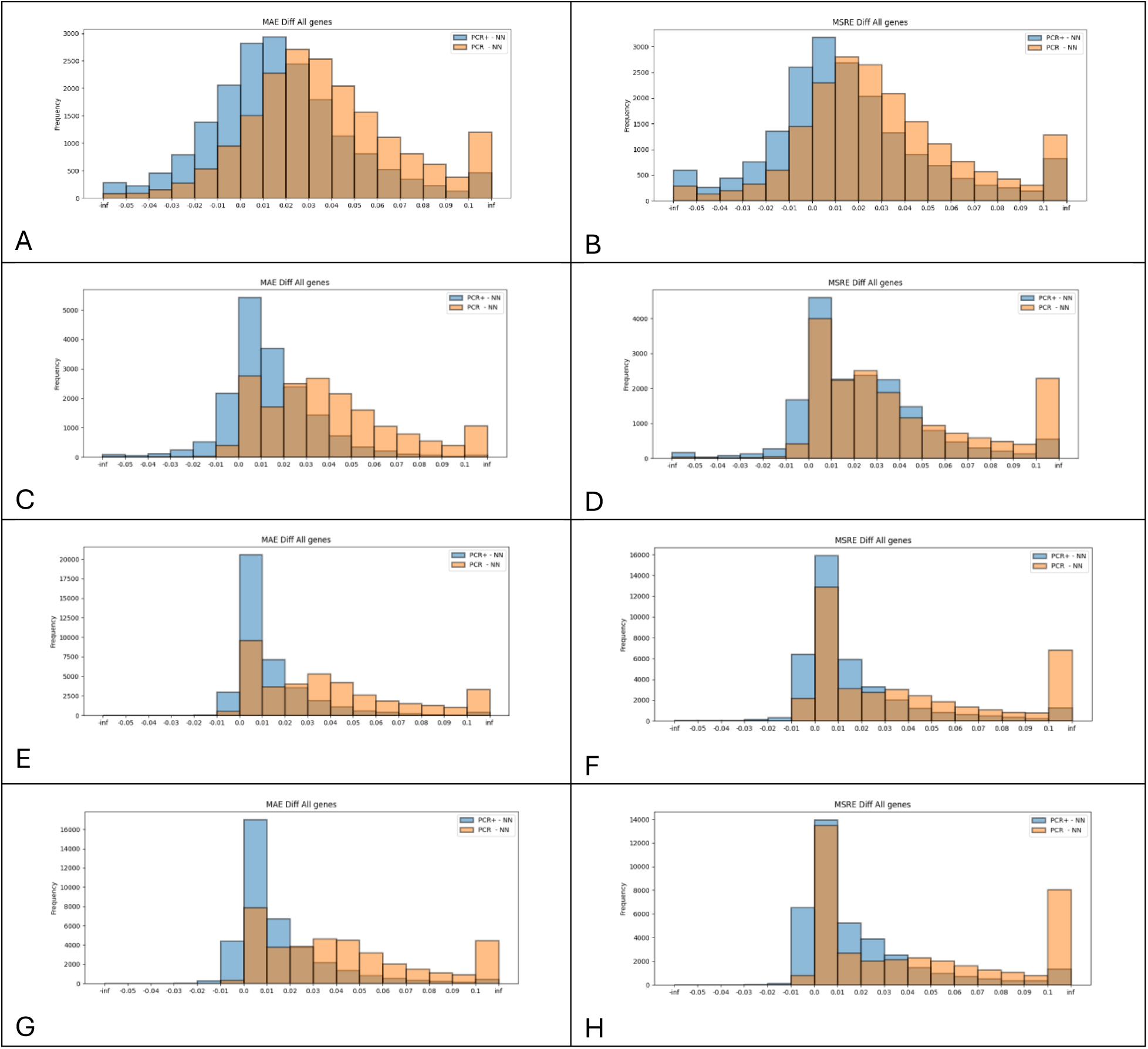
Improvement of NN-MoE and PCR+ methods compared to baseline (PCR) for [A,B] zebrafish (14,924 sample dataset), [C,D] rat (26,716 sample dataset), [E,F] mouse (361,100 sample dataset), and [G,H] human (463,055 sample dataset).

**Table 4:** Median NN-MoE MAE and MSRE for four organisms.

| Organism | Dataset Size | Median MAE | Median MSRE |
| --- | --- | --- | --- |
| Zebrafish | 14,924 | 0.1695 | 0.1478 |
| Rat | 26,716 | 0.1138 | 0.0650 |
| Mouse | 361,100 | 0.0328 | 0.0615 |
| Human | 463,055 | 0.0467 | 0.0855 |

### Developmental Neurotoxicity Case Study

To illustrate the practical impact of transcriptomic extrapolation, we narrowed our focus to data from a recent study [17]. In this study, we conducted a comprehensive evaluation of a subset from a 91-compound library established by the NIEHS Division of Translation Toxicology to support the development of a battery of assays aimed at elucidating pathways underlying developmental neurotoxicity (DNT) following chemical exposure. The 27 selected compounds span diverse chemical classes including PAHs, organophosphates, and insecticides. Using whole transcriptome sequencing of zebrafish embryos exposed to these compounds at concentrations known to later induce developmental malformations, we aimed to capture early transcriptional responses that may underlie the onset of adverse developmental outcomes.

**Figure 6** shows the impact that performing extrapolation of the measured genes can have on the overall ability to detect differentially expressed genes (DEGs). Compared to using measured genes alone, extrapolation using MoE-NN, PCR+ and PCR all consistently improve F1 scores, sometimes dramatically. As expected, most of the improvements are a result of increased sensitivity. At the same time, loss in precision was minimal.

**Figure 6:**
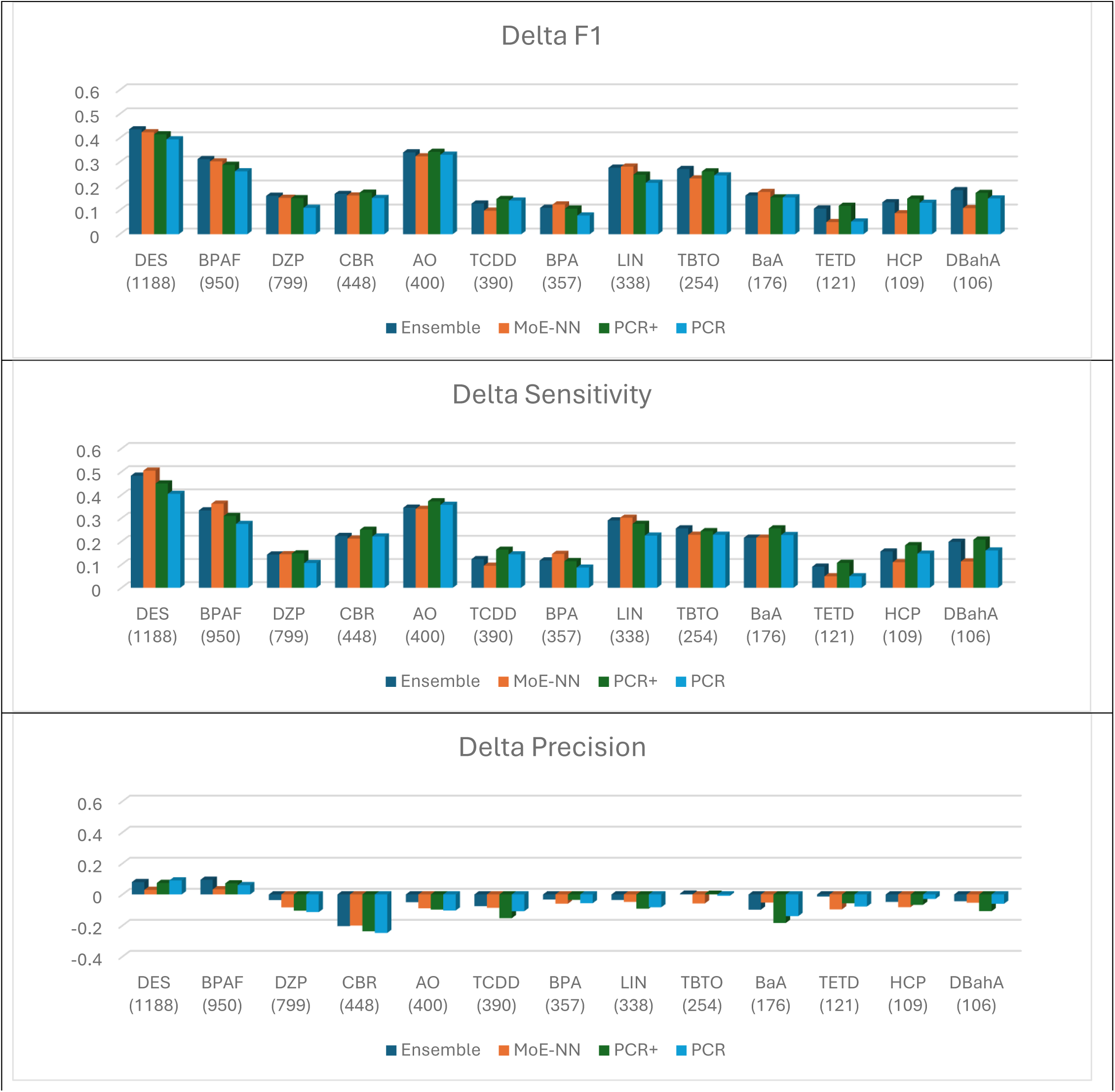
Observed changes in DEG detection performance observed in Developmental Neurotoxicity study for the 10 chemicals with at least 100 DEGs. Comparison is relative to using only measured S1500+ genes. The “Ensemble” method uses the mean of the MoE-NN and PCR+ predictions.

**Figure 7** illustrates the impact that performing extrapolation has on pathway coverage. In the innermost ring we see pathways that are enriched when using only measured genes. Surrounding rings illustrate pathways that are enriched when including extrapolated gene measurements. The outermost ring displays pathways that are enriched using the full original RNA-seq data, which is the gold standard for the data from this study. In most cases, the overall results are consistent across the rings, but the improved pathway coverage and accuracy of gene-level expression arising from extrapolation leads to greater enrichment with a greater signal to noise ratio. Using the TCDD comparison in Figure 7 as an example, extrapolation leads to stronger enrichment (lower adjusted p-values) for HYPOXIA and EPITHELIAL_MESENCHYMAL_TRANSITION, with values much more in line with what is observed in the full RNA-seq data (compared to the results when only measured genes are utilized). For TCDD, COAGULATION, which has an adjusted p-value of 0.0033 (unadjusted p-value=0.0004) in the full RNA-seq data, has a non-significant adjusted p-value of 1 (unadjusted p-value=0.1887) when only the measured genes are used in the pathway analysis. However, once extrapolation is applied, the adjusted p-value for COAGULATION reduces drastically, ranging from 0.0564 to 0.1938 (unadjusted p-value=0.0056 to 0.0271) depending on extrapolation method.

**Figure 7:**
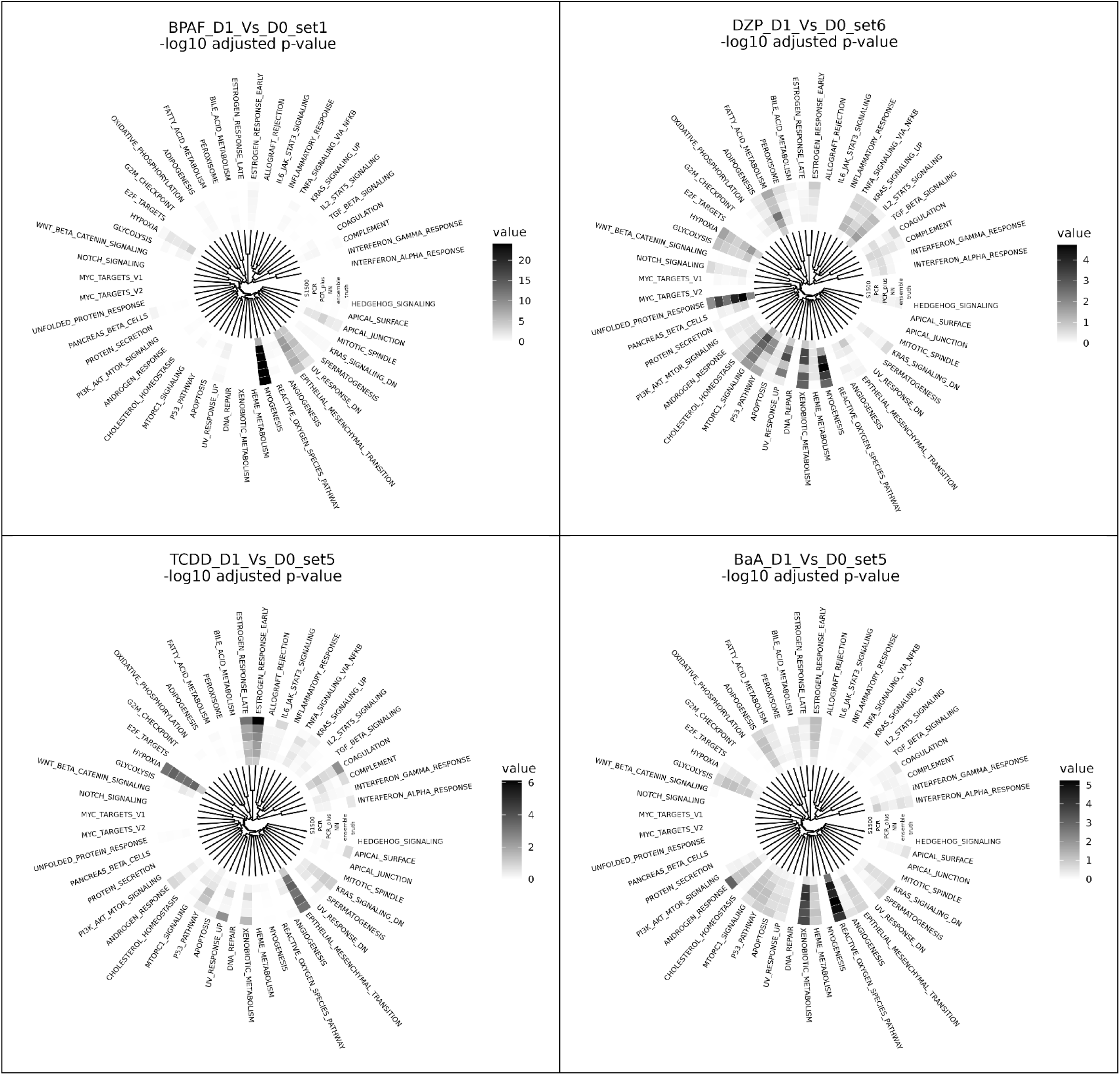
Pathway landscape plots depicting the −log10 adjusted p-values for the significance of Hallmark pathways (n=50) across the analysis levels (measured genes only for innermost ring, respective extrapolated genes included in the DEP analysis for the middle rings, and the true RNA-seq whole transcriptome DEP results in the outermost ring). Pathways are ordered in the circular view based on hierarchal clustering of the gene membership (i.e., gene set similarity). Each compound has a unique scale, with darker shades for the higher −log10 adjusted p-values (ie., lower, more significant, adjusted p-values).

## Discussion

This study extends targeted transcriptome extrapolation strategies to zebrafish, an increasingly important model for developmental toxicology and high-throughput chemical screening. By curating a large and heterogeneous training dataset, we demonstrated that reduced-representation approaches can reliably reconstruct unmeasured gene expression profiles at a fraction of the cost of whole-transcriptome sequencing. Our evaluation of three extrapolation methods—standard PCR, a weighted PCR+ variant, and a neural network mixture-of-experts (NN-MoE)—provides new insights into the performance tradeoffs of alternative modeling approaches in the zebrafish context.

A central finding of this work is the clear impact of biological heterogeneity on extrapolation accuracy. Models trained and tested within the same tissue consistently achieved lower error rates compared to cross-tissue settings. Conversely, substantial performance degradation was observed when models were trained and tested on dissimilar tissues. Exceptions tended to arise between tissues that are anatomically or developmentally related, such as the brain and telencephalon or germ layer–related pairs like liver and intestine. These observations underscore the importance of developmental biology in shaping transcriptomic similarity and suggest that incorporating biological priors, such as germ layer origin or lineage relationships, could further enhance extrapolation strategies.

Our comparison of extrapolation methods revealed distinct advantages of the proposed extensions over the baseline PCR approach. The locally weighted PCR+ method, by tailoring regression to samples most similar to the test set, yielded consistent improvements in both MAE and MSRE. The NN-MoE approach provided the greatest gains, reducing average error by approximately 20% across evaluation metrics. Importantly, the NN-MoE architecture was able to exploit heterogeneity in the training data by dynamically weighting specialized regression heads according to tissue- or study-specific features. This capability is particularly advantageous in zebrafish, where transcriptomic datasets are fragmented across diverse experimental contexts. Furthermore, our application of this method to datasets from several additional species (rat, mouse, and human) suggests that these gains are likely to continue to increase as new high quality zebrafish transcriptomics datasets become available for training.

Despite these advances, several limitations remain. First, while the curated dataset was large by zebrafish standards, it remains modest relative to human or mouse transcriptomic resources. This restricts the diversity of contexts represented in training and likely constrains the generalizability of extrapolated models. Second, although the NN-MoE method demonstrated superior accuracy, it requires greater computational resources to train. Finally, a small subset of genes consistently showed poor extrapolation performance across methods, highlighting inherent limitations in predictability that may relate to unique regulatory dynamics or weak correlation with the Zf S1500+ subset. However, given that we can reliably identify these genes based on their performance on the training data, they can be removed from further analysis, if necessary.

Looking forward, the availability of larger and better-annotated zebrafish transcriptomic datasets will likely enhance extrapolation accuracy and generalizability. Additionally, future work could evaluate cross-species extrapolation approaches, leveraging orthologous gene sets to transfer predictive models between zebrafish, mouse, and human datasets. Such efforts would further expand the translational value of zebrafish extrapolation in toxicology and biomedical research.

## Conclusions

We developed and validated a zebrafish-specific targeted transcriptomics extrapolation framework using the Zf S1500+ gene set and evaluated three modeling strategies. Both the locally weighted PCR+ and NN-MoE methods improved upon the baseline PCR approach, with NN-MoE yielding the most substantial performance gains by dynamically accommodating dataset heterogeneity. Our analyses highlight the critical role of tissue-and lineage-specific context in extrapolation performance and suggest that incorporating developmental biology priors may improve cross-tissue generalizability.

These findings establish a practical and scalable framework for improved transcriptome extrapolation using a deep learning approach in zebrafish, reducing the cost barrier for high-throughput toxicogenomics and enabling broader application of the model in chemical safety assessment. As zebrafish transcriptomic resources continue to grow, we anticipate further gains in accuracy, robustness, and cross-species applicability of extrapolated models, ultimately advancing the integration of zebrafish into predictive toxicology pipelines.

## Acknowledgments

This research was supported in part by the National Institute of Environmental Health Sciences of the National Institutes of Health under Award Number R35 ES031709.

## Notes

### Competing Interest Statement

The authors have declared no competing interest.

